# Efficacy of the entomopathogenic fungus *Beauveria bassiana* in protecting date palm against the red palm weevil *Rhynchophorus ferrugineus* under hot desert climate

**DOI:** 10.1101/2024.07.13.603376

**Authors:** Omer Zer Aviv, Sabina Matveev, Dana Ment

**Affiliations:** Department of Plant Pathology and Weed Research, Agricultural Research Organization, Volcani Center, Rishon LeZion, Israel; The Robert H. Smith Faculty of Agriculture, Food & Environment, The Hebrew University of Jerusalem, Rehovot, Israel

**Keywords:** Red palm weevil, *Beauveria bassiana*, Entomopathogenic fungi, Biological control, Date palm, Seasonal efficacy

## Abstract

The red palm weevil (*Rhynchophorus ferrugineus*; RPW) is an important pest threatening date palm cultivation worldwide. Conventional insecticide-based management is limited by environmental concerns and the development of resistance in RPW populations. Entomopathogenic fungi (EPF), such as *Beauveria bassiana*, offer a promising biocontrol alternative. While laboratory and semifield studies have demonstrated their potential as insecticides, evidence from field-scale applications under diverse environmental conditions remains scarce. This study evaluated the seasonal efficacy and persistence of *B. bassiana* as a preventive treatment against RPW in male and female date palm groves subjected to different cultivation practices in Israel’s Arava region. Healthy palms were equipped with IoTree sensors to monitor RPW infestation dynamics. EPF treatments were applied at the start of each season, and control trees were left untreated. Soil samples were collected seasonally to quantify fungal colony-forming units (CFUs) and assess their association with treatment outcomes. Results demonstrated a significant seasonal influence on EPF efficacy, with autumn treatments yielding the highest reduction in infestation rates, and summer applications showing limited effectiveness. Efficacy also varied between grove types: spring applications were most effective in male groves, whereas autumn treatments were superior in female groves. A positive correlation was found between *B. bassiana* CFUs in the soil and palm health, indicating that EPF persistence enhances treatment success. These findings highlight the importance of seasonal and site-specific optimization of EPF-based biocontrol strategies. Further research is needed to refine application schedules and validate outcomes across larger-scale trials.

## 1. Introduction

The red palm weevil (RPW) *Rhynchophorus ferrugineus* (Olivier) (Dryophthorinae; Curculionidae) is an important pest of palm trees worldwide, particularly the genus *Phoenix*. The RPW life cycle is cryptic, occurring mostly inside the tree trunk (Murphy and Briscoe 1999; Blumberg 2008; Prabhu and Patil 2009; El-Shafie and Faleiro 2020). The female RPW bores holes in the palm tissue with its mouthparts. Within these holes and in natural crevices or wounds in the palm trunk, it lays individual eggs (Ince et al. 2011; Hussain et al. 2013; Matveev et al. 2023), with larvae emerging in 4 to 7 days. The larval stage lasts from 60 to 105 days. Inside the palm trunk, the larva metamorphoses into a prepupal stage that lasts for 3 to 6 days before becoming a pupa, for 13 to 17 days. After this transformation, the adult RPW emerges and lives for 100 to 120 days (Murphy and Briscoe 1999). The RPW is polyphagous and demonstrates rapid dispersion due to its ability to fly long distances in a short period of time (Dembilio and Jaques 2015; Hoddle et al. 2015). Damage to the tree stems from RPW larvae feeding on, and destroying, the palm tissue (Giblin-Davis 2001).

Conventional farming practices often rely on chemical insecticides, such as imidacloprid and thiamethoxam, as the primary means of controlling RPW infestations (Kaakeh 2006; Dhouibi et al. 2017; Chihaoui-Meridja et al. 2019). The insecticides are used as a preventive tool to stop RPW establishment in healthy palms, and as a curative tool in RPW-infested palms. For prevention, imidacloprid and thiamethoxam are applied to the tree’s root area in the irrigation water; for preventive and curative purposes, a mixture of imidacloprid and gamma cyhalothrin is sprayed on the palm’s offshoots and trunk up to 2 m in height. These insecticides have a common mode of action that affects the insects’ nervous system, causing paralysis and death (Kurwadkar et al. 2013). Although these compounds have good efficacy levels, they also have drawbacks, such as the development of resistance to them in RPWs (Wakil et al. 2018) and their potentially lethal toxicity to humans (Mohamed et al. 2009; Hashemi-Domeneh et al. 2016; Mundhe et al. 2017).

Integrated pest management (IPM) is used in many countries, including Israel, to control the spread of RPW (Al-Dosary et al. 2016). The IPM program in Israel is comprised of sanitation, the use of seismic sensors for RPW detection, and preventive and curative treatments (Israeli Extension Service 2022). Sanitation measures include weed removal in the trunk area, the distancing of sprinklers to avoid trunk wetness, and offshoot removal. The latter is performed in the winter when RPW activity is low, and the wound is treated with tree paste containing an asphalt emulsion. Sensors are installed on palm trunks to detect RPW-infested trees (Figure 2 in Mendel et al. 2024a). Another component of IPM is the use of monitoring traps based on the beetle’s aggregation pheromones, which assists in timing the preventive application of pesticides according to RPW adults’ seasonal activity. In organic farming practices, beneficial nematodes are sprayed on the palm’s offshoots and trunk up to 2 m in height as a curative treatment. Laboratory and field trials have proven these nematodes’ pathogenicity to the pest larvae, pupae, and adults (Abbas et al. 2003; Yaacobi et al. 2023). Botanigard (Rimi, Petah Tikva, Israel), a *Beauveria bassiana*-based product, is applied to the same palm parts as a preventive treatment. It is the only preventive treatment registered for organic management practice (Israeli Extension Service 2022).

*Beauveria bassiana* (*Vuliumin*) and *Metarhizium robertsii (Metchnikoff, Sorokin)* (Ascomycota: Hypocreales) infect a wide variety of arthropods and are used commercially in various agricultural crops (Roberts and Hajek 1992; Van Lenteren et al. 2018). These characteristics make them the most suitable entomopathogenic (EPF) species for biological pest control (Faria and Wraight 2007). Commercial EPF insecticides based on *B. bassiana* species are widely used in the United States, the European Union, Brazil, Japan, and Mexico for various crops against numerous pests of different orders (Mascarin and Jaronski 2016). Abiotic factors, such as temperature, ultraviolet (UV) radiation, and humidity, affect EPF’s survival, growth, and virulence, with different isolates exhibiting differential survival based on the climate in the area from which they were isolated. Thus, there can be intraspecies variation in their tolerance to stress. Temperature is considered a major factor affecting EPF growth and virulence. The longevity of ascomycete conidia is optimal at low temperatures, whereas growth and disease progression in hosts are optimal at higher temperatures (Ment et al. 2017). UV is the most influential abiotic factor on the survival of cells in general. For EPF in particular, even the most resilient isolates will not survive after a few hours of direct UV exposure (Fernandes et al. 2015). UVA radiation causes reactive oxygen species accumulation in the cell, while UVB radiation changes the structure of the pyrimidine nucleotides (Griffiths et al. 1998). UVA radiation has been found to reduce the sporulation and endurance of *Metarhizium anisopliae* conidia (Braga et al. 2001), and UVB showed the same effects on *B. bassiana* (Fernandes et al. 2007). Of note, the latter experiment showed that distance from the equator affected the endurance of the UVB-exposed isolates, with those produced near it being more resistant to UVB than those produced further away from it. EPF formulations containing UV-protection additives, such as vegetable oils, adjuvant oils, oil-soluble sunscreens, optical brighteners, and clay, increased the survival of conidia under field conditions (Ment et al. 2017). High relative humidity (RH) is especially important for spore germination and fungal sporulation. In a laboratory experiment, 100% RH ensured rapid and efficient infection of sawtoothed grain beetle by *B. bassiana*. At <90% RH, infection occurred sporadically and at low rates (Searle and Doberski 1984). Whereas high RH is crucial for conidial germination and disease progression in insects in aerial habitats, low RH is a prerequisite for conidial longevity in the environment (Ment et al. 2017, 2020).

Several laboratory studies have shown that EPF can control RPW populations by reducing their egg-hatching rate and eliminating both larval and adult stages, and can complete their life cycle on dead RPW cadavers (Gindin et al. 2006; Al-Keridis et al. 2020). Additional laboratory and semifield experiments have confirmed that species such as *B. bassiana* and *M. anisopliae* can infect and eliminate all life stages of the RPW with high efficacy rates, sometimes achieving mortality rates of up to 100% (Dembilio et al. 2010; Güerri-Agulló et al. 2010; Fong et al. 2018). Recently, Ment et al. (2023) reported that Velifer®, a *B. bassiana* product, and *Metarhizium brunneum* (Mb7) result in 60% to 88% female mortality. Mb7—as a conidial suspension or powder—resulted in 18% to 21% egg-hatching rates, approximately three times less than in the non-treated control. Treating palms under greenhouse conditions with Mb7 significantly decreased signs of infestation and resulted in palm protection. Regarding fungal persistence, a previous study from our research group investigated the persistence of EPF in date palm soil and trunks in Israel: viable fungal units persisted in the soil up to 280 days and in the trunk up to 90 days. In addition, the numbers of EPF colony-forming units (CFUs) in samples taken from the soil and trunk were moderately correlated, with their levels decreasing similarly over time (Ment et al. 2020; Livne 2024). EPF have also been found effective against RPW in several field trials (Sabbahi and Hock 2023). However, the persistence and efficacy of EPF in RPW control have not been tested in large-scale field trials, nor have they been evaluated under different agricultural practices or climate conditions. Another unexplored subject is the relationship between the quantity of EPF propagules present in the soil and the level of RPW infection in palms.

In the present study, we determined EPF survival and efficacy in protecting palms against RPW under different abiotic conditions and agricultural management practices; and analyzed the correlation between palm-protection rate and EPF persistence in the soil. To our knowledge, this is the first field study conducted in palm groves of EPF persistence and efficacy against RPW. The study’s hypotheses are twofold. First, we hypothesized that under abiotic factors that are optimal for fungal survival and establishment, fungal efficacy in protecting palms against RPW will increase. Second, we posited that variations in palm canopy height, and the application of chemical fertilizers vs. compost, will have distinct effects on fungal survival in the soil. We also predicted a positive correlation between the number of EPF colonies (as CFUs) surviving in the soil and the number of healthy treated trees. To test these hypotheses, we chose suitable groves, installed sensors on the palm trees, treated them with EPF-based products, monitored the palms for RPW infestation, and compared the results to untreated palms. Soil samples were collected and EPF persistence was quantified at the beginning and end of each EPF treatment. The experimental period spanned a little over 2 years, with 15 experiments in total.

The aim of this research was to: (i) evaluate the effectiveness of EPF application under various climate conditions in reducing the prevalence of RPW-infested palms in two types of palm groves (tall organic grove vs. short conventional grove); (ii) determine the correlation between EPF persistence and palm protection in treated palms, considering both seasonal variations and differences in morphology and agricultural practices between the two grove types.

## 2. Materials and methods

### 2.1. Date palm grove properties

Date palm groves were selected at Moshav Idan, located in the central Arava Valley in Israel, which is a hot desert area (Fig. 1a, Table 1). The suitability of the groves, as well as each palm tree in the grove, was determined according to specific criteria. The palm groves were 6 to 8 years old when the experiments were initiated (June 2021). RPW infestations had been previously reported in the area of the groves by extension services and local farmers. The two types of groves in which the experiments were carried out were 1.5 km apart (Fig. 1b–c).

**Fig. 1.**
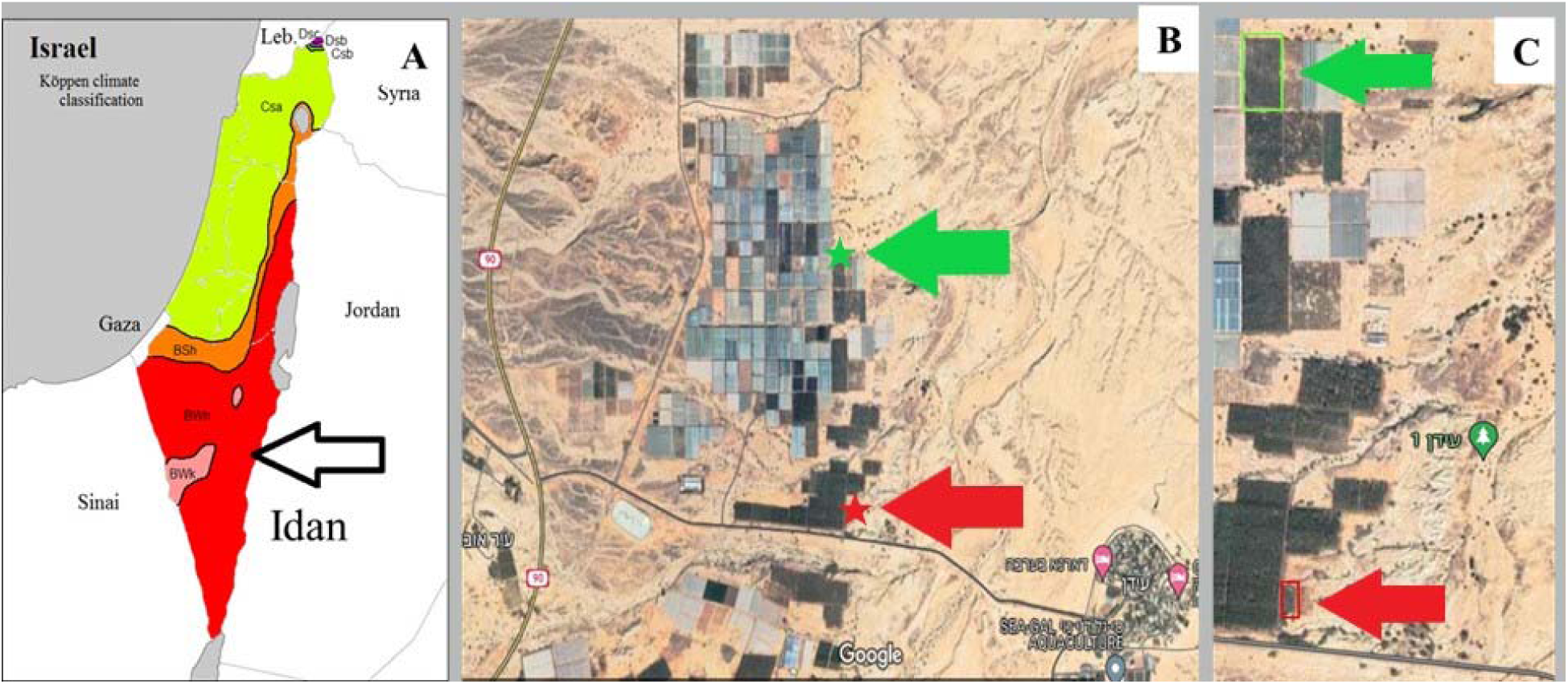
The study area and plantation location in Moshav Idan in the central Arava Valley. (a) Moshav Idan marked on the Koppen classified map of Israel with a climate designated as BWh, Hot desert (Source: Fleischer 2018). (b, c) The studied plantation location is marked in green for Medjool variety palms and red for male palms (Source: Google Maps).

**Table 1.** Descriptive information for the experimental groves.

| Category | Male grove | Female grove |
| --- | --- | --- |
| Total number of palms | 56 | 250 |
| Variety | Mixed varieties | Medjool |
| Agricultural practice | Conventional | Organic |
| Coordinates | 30°48'36.115" N<br>35°16'31.090" E | 30°49'24.611" N<br>35°16'27.808" E |
| Planting year | 2015 | 2014 |
| Fertilization<br>(type, period of application) | Uran 32,<br>March–July | Compost,<br>March |
| Growth retardant application<br>(name, period of application) | Uniconazole,<br>April, every 2 years,<br>from 2022 | - |
| Curative treatment for RPW given<br>(insecticide active compound<br>name/species) | Imidacloprid | Beneficial nematodes<br>(*Steinernema*<br>*carpocapsae*) |
| RPW-infested palms (%)<br>(measured by sensors, 2021) | 13.2 | 3.2 |
| Average trunk height (m)<br>at the beginning of the trials, 2021 | 1.5–2 | 2.5 |

#### 2.1.1. Female grove

The female date palm grove was an organic grove, planted in 2014, comprising 250 female palms of the Medjool variety. The palm fruit are harvested each year in August–October. Fertilization consists of applying compost to the area around the palm trunk in March. Palms are not treated with any growth retardant. Morphology was typical for Medjool variety plantations of similar age, with a uniform trunk height of approximately 2.5 m at the beginning of the experiment. Two weeks before the experiments were initiated, 3.2% of the palms in the grove were considered infested with RPW. Curative treatment measures for RPW using beneficial nematodes were performed before and during the experiments.

#### 2.1.2. Male grove

The male palm grove was planted in 2015 and comprised 56 male palms of mixed varieties. Spathes are harvested every year in February–April to produce date palm pollen. Chemical fertilization is performed every year from March–July with Uran 32 (16% urea, 8% ammonium, and 8% nitrate, v/v). Palms are treated with the growth retardant uniconazole every 2 years to slow vegetative growth, as reflected by lower trunk heights than those in untreated palms of similar age, and shorter leaves which bend slightly toward the ground. Trunk height was 1.5–2 m at the beginning of the experiments. This grove demonstrated a higher infestation rate prior to our trials than the female grove, with 13.2% of its palms considered infested with RPW according to acoustic sensors 2 weeks before experiment initiation. Before and during our study trials, RPW-infested palms were treated with imidacloprid drench as a curative measure. However, these treated palms were not categorized as either control or EPF-treated palms in our study; EPF-treated and control palms were treated with beneficial nematodes.

### 2.2. Experimental procedures

Each tested palm was equipped with a transmitting IoT seismic sensor (Agrint Rockville, MD, USA), indicating its health index status, for at least 1 month prior to initiating the experiments. The palms in each grove were divided into three groups: Control (untreated), Treatment 1 (one type of EPF treatment), and Treatment 2 (another type of EPF treatment); the main treatment applied during the study was spray application of Botanigard (LAM International, distributed in Israel by Rimi). The palms of each experimental group were arranged in clusters (Fig. 2). Control and treatment groups were irrigated on the same watering schedule and were treated equally preharvest and postharvest according to each grove’s regimen. We sprayed each treatment group with either Velifer (BASF, South Africa) or Botanigard emulsion in experiments 1–4 and 10–12. Afterwards, in experiments 5 and 6 and 13 and 14, the Velifer-treated groups received Botanigard + the surfactant Silwet L-77 (L77) as emulsifier. Subsequently, these treatments were also replaced in experiments 7–9 and 15 with a double concentration of Botanigard emulsion (Botanigard X2) (see Table 2 for a description of all experiments, which will be further described in section 2.3).

**Fig. 2.**
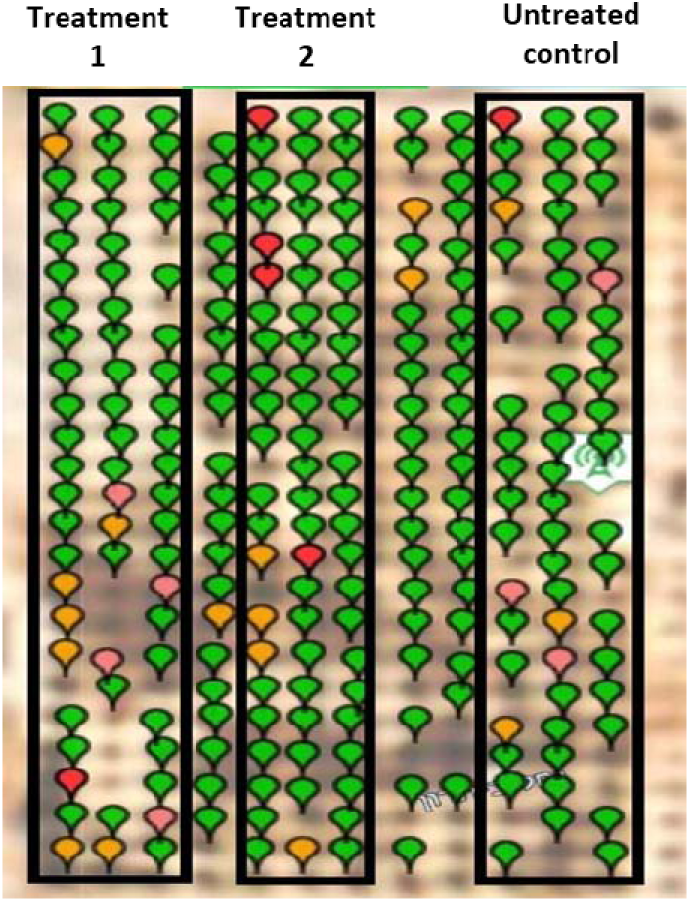
Satellite image of the female grove. Each map pin represents a sensor installed on a palm. Green, clean; yellow, suspect; red, RPW-infested. In this cluster arrangement, each group was separated by 1 or 2 rows of palms. (Source: Agrint Sensing Solutions 2021, website screenshot).

**Table 2.**
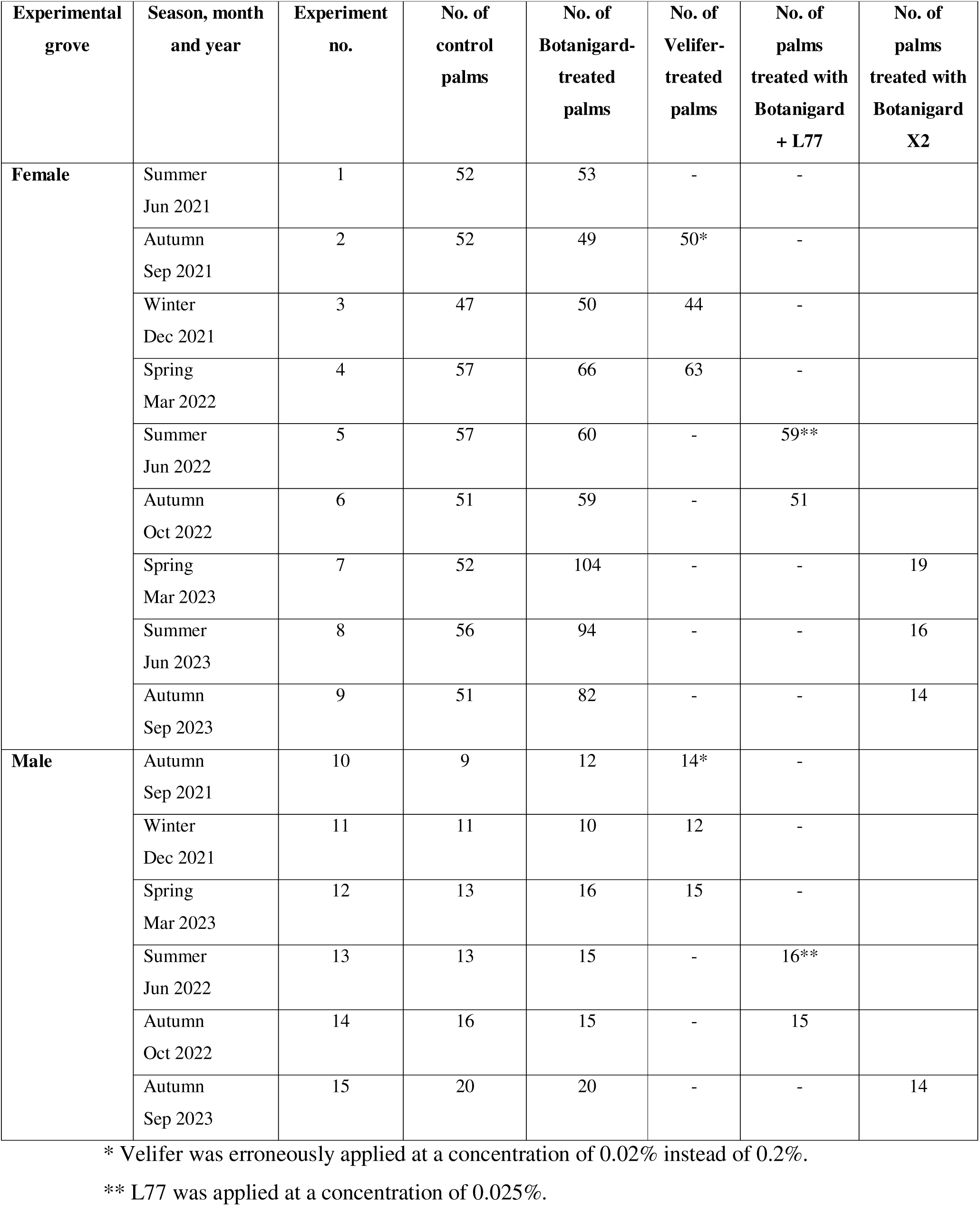
Description of the experiments in the Arava according to name of site, season, and number of palms in each group.

### 2.3. EPF application

Each palm in an experimental group was sprayed manually using a spray gun attached to a tractor-mounted hydraulic sprayer with a capacity of 500 L. To achieve 0.2% (v/v) Botanigard emulsion, 1 L of Botanigard emulsion was diluted with 499 L of fresh water. Before each application, the spray gun flow rate was measured, and spraying time was calculated to apply 8–12 L per palm. The Botanigard + L77 group was sprayed with 0.2% Botanigard emulsion mixed with water and 0.025% or 0.1% (v/v) L77 spreader according to the trial protocol (Table 2). Application was performed by rotary spraying of each palm trunk from its bottom to a height of 1.5 m, and the ground surrounding the palms in a radius of 0.5 m from the trunk base. The treatment was applied once at the beginning of each experiment. EPF preparation specifications are given in Table 3. Same application practices were performed for Velifer as well.

**Table 3.** Commercial EPF product information.

| Commercial name | Fungus | Isolate | Manufacturer | Formulation | Application method | Dose |
| --- | --- | --- | --- | --- | --- | --- |
| Botanigard | *Beauveria bassiana* | GHA | LAM International | Emulsion | Spraying | 0.2%<br>v/v |
| Velifer | *Beauveria bassiana* | PPRI 5539 | BASF | Emulsion | Spraying | 0.2%<br>v/v |

### 2.4. Climate data

The climate databases for the study area were collected from the archives of the Northern Arava Research and Development Center. Monthly mean, maximum, and minimum temperatures at 2 m (°C), RH (%), solar radiation intensity (W/m²), and monthly mean reference evapotranspiration (ET_0_) determined by FAO 56 Penman Monteith equation (Allan et al. 1998) were recorded for each month during the experiments. The monthly averages for these factors were determined using hourly measurements and a standard time base of 24 h. The weather station is located approximately 7 km southwest of our field trial location.

### 2.5. Sensors for RPW activity

Agrint IoTree sensors (Agrint Sensing Solutions) were used to detect RPW activity inside the palm trees. Each palm had a sensor installed on its trunk at a height of 1 m from the ground (Fig. 2). After a calibration period of 2 weeks, the sensor was activated. The sensor’s mechanism of action is based on its ability to recognize the specific vibration of RPW larvae in the palm’s trunk. Each sensor’s vibration information is transferred to a hub, which sends it to a server in real time. The vibration duration and frequency are processed and compared to the vibrations from other palms. After processing, the palm is categorized as clean, suspect, or infested with RPW. The information is shown on Agrint’s website or mobile application. Palms are numbered and displayed on a satellite map with an indication of their infestation levels. Palms marked in green are considered clean, those marked yellow are suspect, and those marked in red are considered infested with RPW (Fig. 2).

### 2.6. Fungal survival evaluation

#### 2.6.1. Soil sample collection for quantification of fungal propagule survival

Soil samples were collected from each of the groves on the day of EPF application, before and after the treatment. Before each application, five samples were collected from each group in the male grove, and ten samples were taken from each group in the female grove. Following the application, two samples were taken from each group treated with EPF in the male grove, and three samples were taken from each EPF-treated group in the female grove. Five different spots around the palm trunk were sampled with a trowel. Samples taken prior to application served to examine EPF survival rates from the previous application. Post-application samples were used to determine the initial number of EPF CFUs and validate emulsion vitality. The difference in CFU quantity counted immediately after application and on the last day of the experiment (before the re-application of the consecutive experiment) served to measure CFU loss during each experiment.

#### 2.6.2. Soil moisture-content determination

Each sample of palm soil was well-mixed, and approximately 10 g of the wet sample was extracted, poured, and placed on a 90-mm Petri dish labeled with the number and weight of the soil sample. The plate was left to dehydrate for 1 week in a dark room, then weighed again to determine its final weight, for moisture-content determination. The moisture-content percentage was calculated as follows:

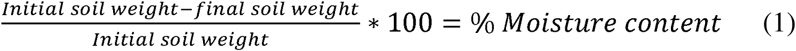

#### 2.6.3. Quantification of CFUs in dry soil

Using the samples taken for moisture-content determination, we also conducted a quantitative analysis of CFU/g dry soil. First, 1 g of wet soil from each sample was mixed with 10 mL sterile water containing 0.01% (v/v) Triton X-100 surfactant for better extraction. The sample was transferred to a rotating incubator for 2 h at 25 °C and 200 rpm. Finally, three 50-µL replicates of the suspension were uniformly sown on 90-mm Petri dishes that contained fungus-selective Sabouraud Dextrose Agar (SDA) supplemented with 1 µL/1 mL chloramphenicol and 1 µL/2 mL dodine. Sown samples were incubated in a dark room at 28 °C for 5–7 days according to colony appearance and density. CFUs were then counted and multiplied by 200 to compensate for the dilution factor. To calculate the weight of the dry soil, each sample’s retrieved moisture content was subtracted as follows:

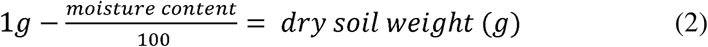

Finally, the normalized CFU count was divided by the normalized dry soil weight to give the quantity of CFU/g dry soil.

### 2.7. Statistical analyses

All statistical analyses were conducted using JMP^®^ software version pro-16 (SAS Institute). The number of infested and healthy palms and the CFU survival data for each group were recorded using spreadsheet software developed by Microsoft Excel. Soil sample CFU counts were repeated three times to obtain representative data. A fixed significance level (α) of 0.05 was used for all tests.

#### 2.7.1. EPF palm-protection efficacy test

To test whether EPF treatments had a significant effect as a protective measure against RPW in each season, we used Chi-square (χ^2^) or Fisher exact tests. At the end of each season, infested and healthy palms were counted in each group; they were categorized separately and their frequency registered with respect to total number of palms in the group. In the next stage, a Fit Y by X function was used to test the relationship between the treatment and the frequency of infested and healthy palms. Treatment or season was set as the X factor, accordance to the type of analysis, and palm health was set as the Y factor. Frequency of each group’s healthy or infested palms was set as the Frequency value. Where 20% or more of the cells in the χ^2^ test table had an expected count of less than 5, Fisher’s exact test was used. To compare the efficacy rates of EPF preparations in each season, healthy palm frequencies in the different EPF treatments were compared to each other without including the control group. To test the seasonal variations in efficacy of each EPF preparation, the rate of healthy palms in each treatment was compared to that in the control group.

#### 2.7.2. Effect of climate variables on EPF treatment efficacy

The ratio of healthy to total palms was recorded in the control and EPF groups each month, as described in section 2.7.1. To avoid distortions in sample sizes, the number of infested palms each month was subtracted from the group’s total number of palms in the following month, until the next EPF application. For example, if 2 out of 52 palms were infested with RPW in the control group in January, in February, those 2 palms were removed from the total number of palms in that group, and the healthy-to-total palm ratio was determined from the corrected total number of 50 palms, for that group. After summing the healthy-to-total palm ratios for each month, efficacy of EPF treatments was determined in relation to the control group, using Henderson–Tilton’s formula. Originally, this formula was used to calculate the efficacy of insecticides with unequal sample sizes in the treated vs. control groups (Henderson and Tilton 1955). We modified it to calculate the protection efficacy of EPF vs. the control treatment as follows:

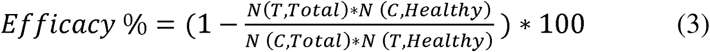

where:

N (T, Total) = total number of treated palms

N (C, Total) = total number of control palms

N (T, Healthy) = number of palms that remained healthy in the treatment group

N (C, Healthy) = number of palms that remained healthy in the control group.

The formula’s results were presented as quantitative percentages, where a higher ratio of healthy palms among EPF treatment groups vs. the control group yields a higher positive value, a lower ratio yields a lower negative value, and identical efficacy rates in the two groups results in 0%. Statistical analysis was performed using the Fit Y by X function, where monthly ET_0_ values were set as the affecting variable (X), and monthly calculated efficacy percentages were set as the dependent variable (Y). The regression between these variables was examined in each grove separately. Due to multicollinearity between temperature, RH, and UV, ET_0_ was chosen as the index, increasing with higher UV and temperature, and lower RH.

#### 2.7.3. Seasonal variations in EPF survival rate in female and male groves

EPF survival rates were evaluated for each grove in different seasons. CFU loss per day was calculated for each sampled palm as:

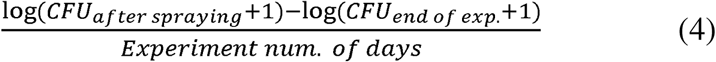

CFU/g dry soil numbers were registered immediately after spraying and at the end of the experiment for each sampled palm. A logarithm was applied with each sample to normalize data distribution and stabilize the variance across the different samples. Prior to log application, 1 CFU was added for each sample to avoid zero values in the data. To normalize the time factor, CFU loss per day was calculated. The log-transformed values within each group were calculated as follows: the final value at the end of the experiment was subtracted from the initial value recorded at the beginning of the experiment, and this difference was then divided by the number of experimental days. The calculated data for each experiment were recorded on an Excel sheet and each sample was categorized by grove and season. CFU loss per day was defined as the dependent factor. Before each ANOVA test, all dependent factors were subjected to Levene’s test to assess equality of the variances (α < 0.05).

To determine the impact of season and grove on EPF survival, whole-model analysis was carried out using the fit model option in JMP. Season, grove, and their combination were defined as the contributing factors, using the cross option. Rank transformation was applied to CFU loss per day values to achieve equal variances. A post-hoc Tukey-HSD test was conducted to examine differences in survival between tested seasons.

To determine the impact of season on EPF survival in each grove, each season was considered the independent variable (X factor) for each grove separately. When equal variances were observed, one-way ANOVA was performed. In the case of unequal variances, Welch’s t-test was utilized to assess the seasonal variances. Where significant variations were found between seasons, multiple comparisons were performed between survival rates for each pair of seasons. Each comparison was subjected to another Levene’s test to check for homogeneity. If variances were equal, a pooled t-test applied; otherwise, t-test was used. Bonferroni correction was applied to adjust the level of significance by dividing the significance level by the number of comparisons. Hence, the adjusted α level was set to: 0.05/3 = 0.0167.

Differences in EPF survival rate between groves were determined in each season, with grove as the independent factor. Season comparisons with equal variances in survival rates were subjected to pooled t-test, while unequal variances were analyzed by Welch’s t-test.

#### 2.7.4. Association between EPF survival and palm health

To examine the relationship between the quantity of EPF found in the soil and the relative number of healthy palms in each group, χ^2^ test was carried out. The relationship was tested with the Fit Y by X function. In each grove, the average CFU count was calculated for each EPF and control group, using the samples collected from each group at the end of each season, prior to the next application of EPF. As described in section 2.7.3, 1 CFU was added to each sample and a logarithm was applied. The calculated log CFU average at the end of the season was defined as the X factor, and palm health in the group was defined as the Y factor. Palm health and frequency were calculated as in section 2.7.1. To create a graph showing the relationship between two variables, a probability formula based on the χ^2^ results was created and saved on the data sheet. Then, the graph builder function of JMP^®^ software was used, where mean CFU survival was defined as the X factor and the calculated probability for healthy palms was defined as the Y factor.

## 3. Results

### 3.1. Climate variables

Table 4 presents the summary of mean, maximum, and minimum temperature, RH, and UV radiation across the experiment’s different seasons. Detailed data on climate conditions for each month are provided in Table S1.

**Table 4.** Summary of measured climate data during the study by season.

| Season | Temperature<br>(°C) |  |  | Relative humidity<br>(%) |  |  | UV radiation<br>(W/m<sup>2</sup>) |  |  | ET<sub>0</sub><br>(mm/day) |
| --- | --- | --- | --- | --- | --- | --- | --- | --- | --- | --- |
|  | Mean | Max | Min | Mean | Max | Min | Mean | Max | Min | Mean |
| Summer | 32.9 | 46.2 | 21.5 | 39 | 74 | 6 | 312 | 1026 | 0 | 7.35 |
| Autumn | 23.6 | 40.9 | 6.8 | 51 | 96 | 13 | 171 | 921 | 0 | 3.12 |
| Winter | 15.8 | 26.9 | 3.1 | 53 | 92 | 15 | 126 | 825 | 0 | 1.92 |
| Spring | 23.7 | 42.4 | 6.5 | 39 | 92 | 6 | 247 | 1037 | 0 | 4.68 |

### 3.2. Palm-protection efficacy of EPF treatments under varying climate conditions and agricultural practices

#### 3.2.1. Seasonal effect on protection efficacy of EPF treatments

None of the experiments conducted in the female date palm grove during the summer seasons showed significant results for EPF treatment protection against RPW infestation (Table 5). In the autumn seasons in this grove, EPF protection showed greater efficacy (Table 5). In the single winter experiment performed in this grove, none of the palms in the treatment or control groups were infested with RPW, making it impossible to measure the effectiveness of EPF protection (Table 5). In the two spring experiments in this grove, EPF protection was not significant compared to control.

**Table 5.** Summary of experiments by grove, period of application, experiment number, total number of palms in a group, and the percentages of those palms that remained healthy.

| Grove | Season and month of application | Exp. no. | Control<br>No. of palms<br>(Healthy %) | Botanigard<br>No. of palms<br>(Healthy %) | Velifer<br>No. of palms<br>(Healthy %) | Botanigard +<br>L77<br>No. of palms<br>(Healthy %) | Botanigard<br>X2<br>No. of palms<br>(Healthy %) | P-value |
| --- | --- | --- | --- | --- | --- | --- | --- | --- |
| **Females** | Summer<br>Jun. 2021 | 1 | 52<br>(100%) | 53<br>(96%) | - | - | - | 0.4952<br>@ |
|  | Autumn<br>Sep. 2021 | 2 | 52<br>(92%) | 50<br>(100%) | - | - | - | 0.1179<br>@ |
|  | Winter<br>Dec. 2021 | 3 | 47<br>(100%) | 50<br>(100%) | 44<br>(100%) | - | - | - |
|  | Spring<br>Mar. 2022 | 4 | 55<br>(95%) | 64<br>(91%) | 61<br>(93%) | - | - | 0.7674<br>@ |
|  | Summer<br>Jun. 2022 | 5 | 54<br>(91%) | 59<br>(90%) | - | 55*<br>(84%) | - | 0.4697<br># |
|  | Autumn<br>Oct. 2022 | 6 | 51<br>(92%) | 59<br>(97%) | - | 49<br>(98%) | - | 0.3879<br>@ |
|  | Spring<br>Mar. 2023 | 7 | 52<br>(98%) | 104<br>(92%) | - | - | 19<br>(84%) | 0.1049<br>@ |
|  | Summer<br>Jun. 2023 | 8 | 56<br>(95%) | 91<br>(88%) | - | - | 16<br>(82%) | 0.2141<br>@ |
|  | Autumn<br>Sep. 2023 | 9 | 51<br>(88%) | 82<br>(93%) | - | - | 14<br>(93%) | 0.8206<br>@ |
| **Males** | Autumn<br>Sep. 2021 | 10 | 9<br>(67%) | 12<br>(92%) | - | - | - | 0.2722<br>@ |
|  | Winter<br>Dec. 2021 | 11 | 11<br>(82%) | 10<br>(80%) | 12<br>(92%) | - | - | 0.7094<br>@ |
|  | Spring<br>Mar. 2022 | 12 | 12<br>(75%) | 16<br>(100%) | 13<br>(100%) | - | - | **0.0223**<br>@ |
|  | Summer<br>Jun. 2022 | 13 | 10<br>(100%) | 16<br>(87%) | - | 15*<br>(93%) | - | 0.7749<br>@ |
|  | Autumn<br>Oct. 2022 | 14 | 16<br>(87.5%) | 15<br>(80%) | - | 13<br>(85%) | - | 0.8803<br>@ |
|  | Autumn<br>Sep. 2023 | 15 | 20<br>(85%) | 20<br>(100%) | - | - | 14<br>(93%) | 0.1808<br># |

\*L77 was applied at concentration of 0.025%.
#Calculated by $\chi^2$ test.

Although two summer experiments were conducted in the male palm grove, that in 2023 could not be analyzed due to sensor malfunction, leaving only experiment 13 in 2022 (Table 5). Three autumn experiments were conducted in this grove, showing that protection was not significant. The winter experiment in that grove showed no significant protection.

The spring experiment conducted in the male grove in 2022 indicated a significant protection rate (Table 5). The spring experiment of 2023 was not analyzed due to sensor malfunction.

#### 3.2.2. Effects of treatment and season on protection efficacy of EPF

Comparison of protection efficacy among EPF treatments per season in each grove did not reveal any significant results (Table 6), whereas season had a significant effect on infestation rate among treatment groups compared to the controls (Table 7). None of the EPF treatments resulted in superior protection over the others (Table 6), and seasonal variations did not significantly affect any specific treatment (Table 7).

**Table 6.**
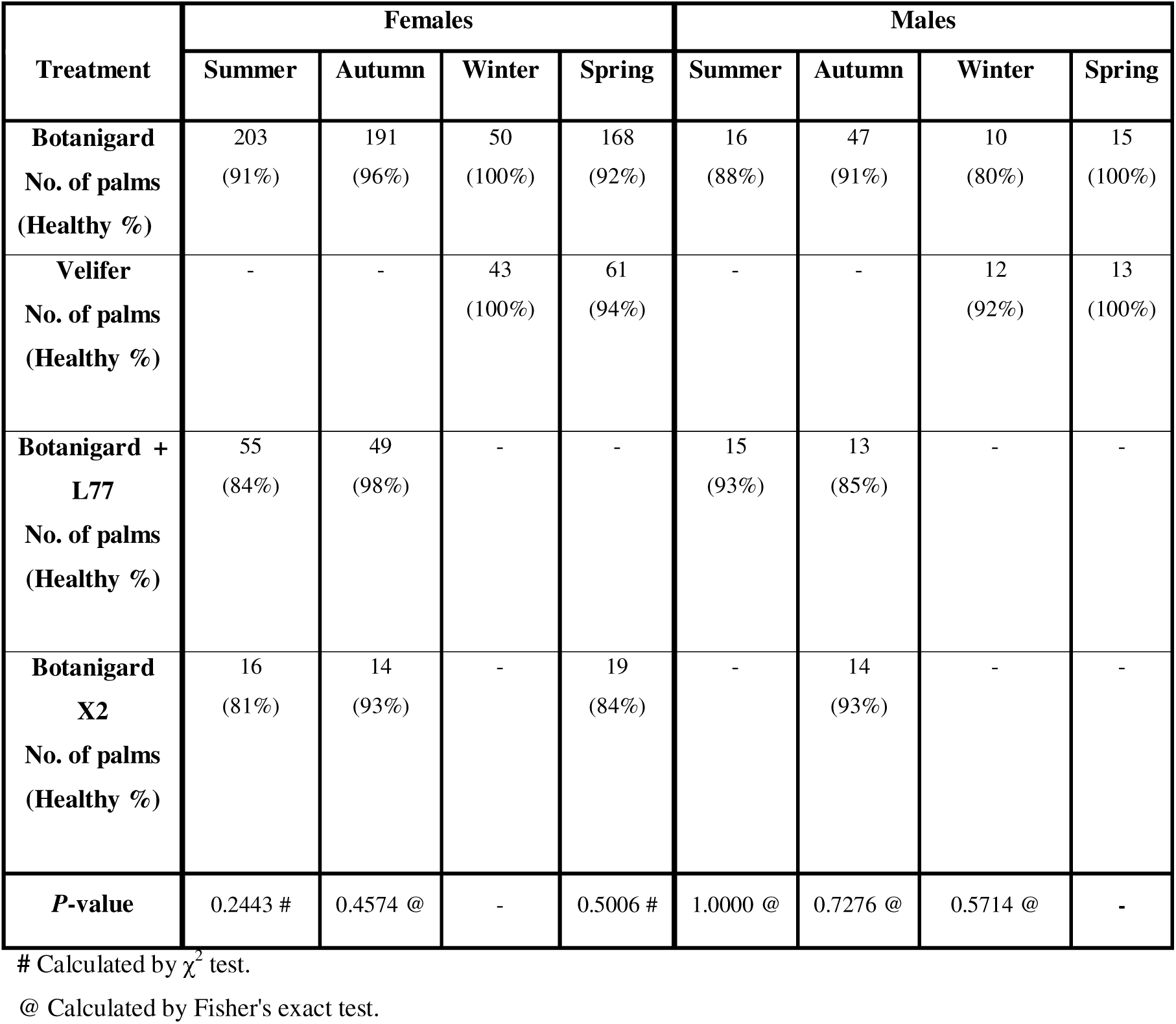
Comparison of EPF treatments’ protection efficacy by season in female and male groves.

**Table 7.** Seasonal impact on EPF treatment efficacy in the female and male groves compared to control groups. #Calculated by χ^2^ test. @ Calculated by Fisher’s exact test.

| Female grove |  |  |  |  |  |
| --- | --- | --- | --- | --- | --- |
| Season | Control<br>No. of palms<br>Healthy % | Botanigard<br>No. of palms<br>Healthy % *P*-value | Velifer<br>No. of palms<br>Healthy % *P*-value | Botanigard + L77<br>No. of palms<br>Healthy % *P*-value | Botanigard X<br>No. of palms<br>Healthy % *P*-val |
| Summer | 162<br>94% | 203<br>91%<br>0.1226 # |  | 55<br>84%<br>**0.0207 @** | 16<br>81%<br>0.0795 @ |
| Autumn | 154<br>91% | 191<br>96%<br>0.0643 # | - | 49<br>98%<br>0.1246 @ | 14<br>93%<br>1.0000 @ |
| Winter | 48<br>100% | 50<br>100%<br>- | 43<br>100%<br>- | - | - |
| Spring | 107<br>96% | 168<br>92%<br>0.1192 # | 61<br>96%<br>0.4632 @ | - | 19<br>84%<br>0.0692 @ |
| Male grove |  |  |  |  |  |
| Season | Control<br>No. of palms<br>Healthy % | Botanigard<br>No. of palms<br>Healthy % *P*-value | Velifer<br>No. of palms<br>Healthy % *P*-value | Botanigard + L77<br>No. of palms<br>Healthy % *P*-value | Botanigard X<br>No. of palms<br>Healthy % *P*-val |
| Summer | 10<br>100% | 16<br>88%<br>0.5077 | - | 15<br>93%<br>1.0000 @ | - |
| Autumn | 45<br>82% | 47<br>91%<br>0.1839 | - | 13<br>85%<br>1.0000 @ | 14<br>93%<br>0.6714 @ |
| Winter | 11<br>82% | 10<br>80%<br>1.0000 | 12<br>92%<br>0.5901 | - | - |
| Spring | 12<br>75% | 15<br>100%<br>0.0752 @ | 13<br>100%<br>0.0957 @ | - | - |

In the female grove, Botanigard was the most effective treatment in the summer; in autumn, Botanigard + L77 was most effective, and in spring, the Velifer treatment. During the winter, both Botanigard- and Velifer-treated palms remained healthy (Table 6). The efficacy rates of Botanigard treatments varied across different seasons, albeit not significantly so. The effectiveness of Botanigard in protecting palms compared to the control showed seasonal variation, with its highest efficacy observed in autumn, and its lowest efficacy in the spring. Notably, in winter, none of the palms in any of the groups were infested with RPW. In spring, the Velifer and control groups had the same percentages of healthy palms (Table 7). Botanigard + L77 treatments also demonstrated significant differences in efficacy across seasons: 98% healthy palms in autumn vs. 84% in the summer. Variations in Botanigard X2 treatment efficacy also varied, albeit not significantly, being most effective in autumn and least effective in the spring (Table 7).

In the male grove, in summer, the Botanigard + L77 treatment was more effective than Botanigard alone; in autumn, Botanigard X2 and Botanigard treatments showed similar rates of efficacy, while the Botanigard + L77 treatment was less efficient; in the winter experiments, Velifer treatment showed higher efficacy than Botanigard, and both of these treatments resulted in 100% healthy palms in the spring (Table 6). In the comparison with the control group, Botanigard treatment efficacy varied over the seasons, from 100% in spring compared to 75% in the control to 80% in the winter vs. 82% healthy palms in the control group. The efficacy of Velifer and Botanigard + L77 treatments compared to controls also showed large variations over the seasons (Table 7).

#### 3.2.3. Combined seasonal effect on palm protection efficacy of EPF treatment

Prevention of RPW infestation by EPF treatments, regardless of the type of EPF preparation, revealed different seasonal efficacy rates among the female and male groves (Table 8).

**Table 8.**
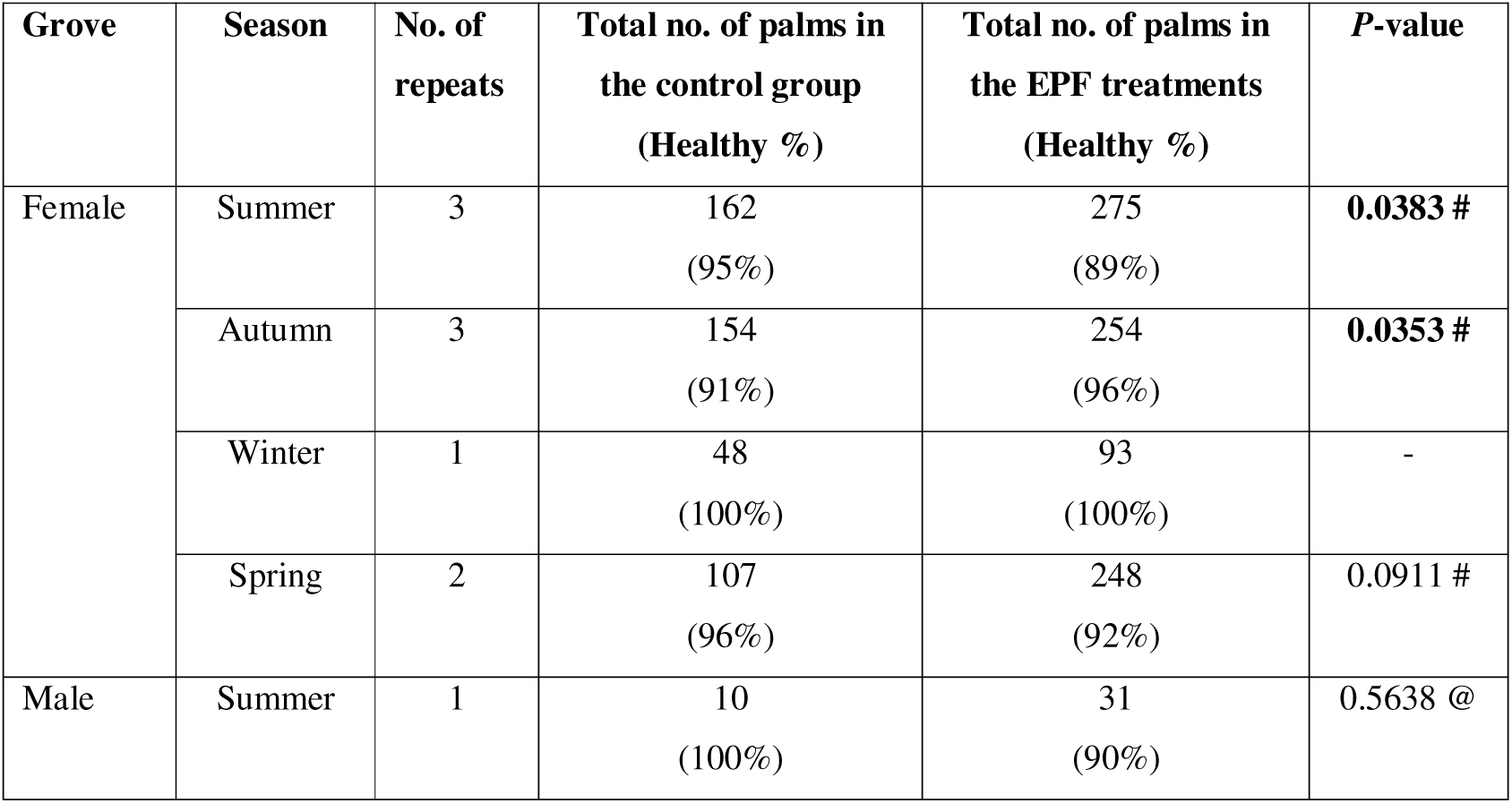
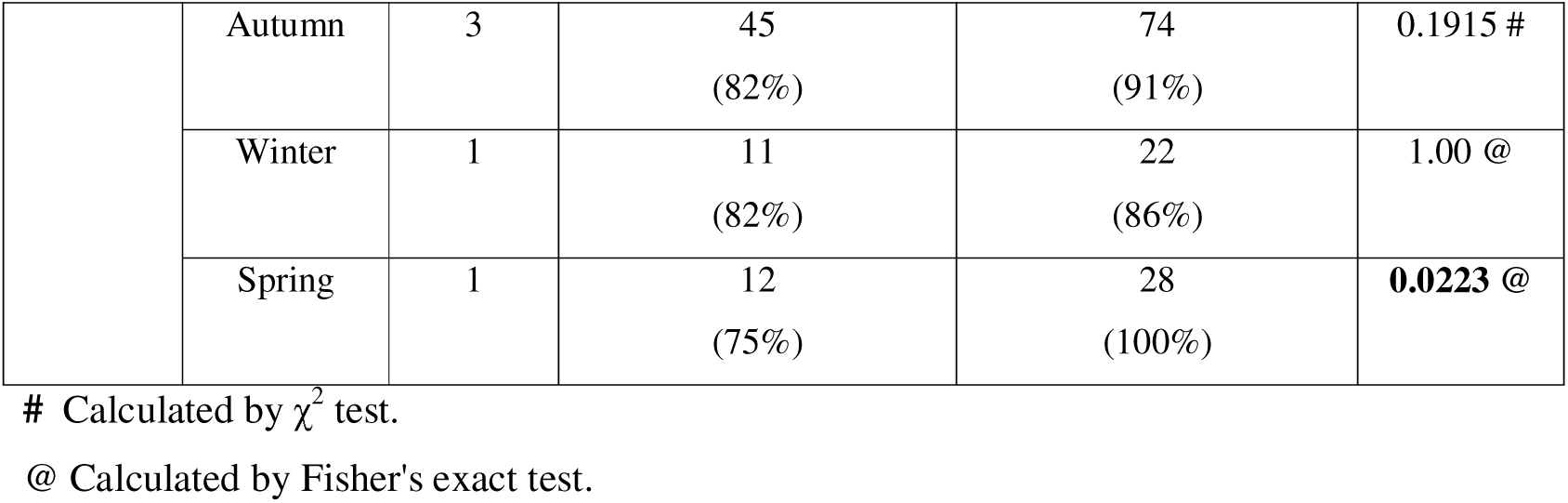
Seasonal comparison of experiments in each grove: number of palms and healthy percent in control and combined EPF treatments and comparison test *P*-values.

| Grove | Season | No. of repeats | Total no. of palms in the control group (Healthy %) | Total no. of palms in the EPF treatments (Healthy %) | *P*-value |
| --- | --- | --- | --- | --- | --- |
| Female | Summer | 3 | 162<br>(95%) | 275<br>(89%) | **0.0383 #** |
|  | Autumn | 3 | 154<br>(91%) | 254<br>(96%) | **0.0353 #** |
|  | Winter | 1 | 48<br>(100%) | 93<br>(100%) | - |
|  | Spring | 2 | 107<br>(96%) | 248<br>(92%) | 0.0911 # |
| Male | Summer | 1 | 10<br>(100%) | 31<br>(90%) | 0.5638 @ |
|  | Autumn | 3 | 45<br>(82%) | 74<br>(91%) | 0.1915 # |
|  | Winter | 1 | 11<br>(82%) | 22<br>(86%) | 1.00 @ |
|  | Spring | 1 | 12<br>(75%) | 28<br>(100%) | **0.0223 @** |

Examination of all three summer experiments in the female grove revealed a significantly higher infestation rate among EPF-treated palms compared to the control group, and a similar, albeit non-significant, trend was observed in the male grove summer experiment (Table 8). The three autumn experiments in the female and male groves showed EPF treatments’ protection of palms from RPW infestation; protection was significant compared to controls in the former, non-significant in the latter (Table 8). For both farming practices (groves), EPF treatments were ineffective compared to control palms during the winter. It is worth mentioning that the overall infestation rate in the winter was relatively low. During the spring, EPF treatments were effective compared to the control group in the male grove but the opposite trend was observed in the female grove, albeit without statistical significance (Table 8).

#### 3.2.4. General protection efficacy of EPF treatments in different seasons

As demonstrated in Table 9, when examining the results in the Arava without considering the effect of agricultural practice or type of EPF preparation, EPF application was only effective in protecting palms from RPW in the autumn season. In the summer, EPF showed inferior palm protection compared to the control group. The winter results showed relatively low and similar infestation rates and in spring, similar healthy palm percentages were observed in both groups.

**Table 9.** General comparison of rate and percentage of healthy palms in control vs. combined EPF groups in each season, with statistical test and *P*-value.

| Season | No. of palms in control group<br>(Healthy %) | No. of palms in EPF treatments<br>(Healthy %) | *P*-value |
| --- | --- | --- | --- |
| Autumn | 199<br>(89%) | 328<br>(95%) | **0.0142 #** |
| Summer | 172<br>(95%) | 306<br>(89%) | **0.0250 #** |
| Winter | 59<br>(97%) | 115<br>(97%) | 1.00 @ |
| Spring | 119<br>(94%) | 276<br>(92%) | 0.5329 # |

### 3.3. Environmental factor effects on EPF treatment efficacy

The effect of environmental factors, as expressed by the ET_0_ value, on the efficacy of EPF treatments was evaluated for male and female groves as described in section 2.7.2. The EPF treatments’ effectiveness was significantly reduced by environmental factors in the female grove, whereas the opposite, albeit non-significant, effect was observed in the male grove (Fig. 3). In the female grove, EPF treatment efficacy strongly decreased with increasing ET_0_ values. Lower ET_0_ values, up to 4, were associated with positive efficacy rates. However, at an ET_0_ of around 4, a turning point occurred where EPF treatments became ineffective and were even less efficient compared to the control group as ET_0_ values increased further. In the male grove, a positive trend was observed between climate variables and EPF efficacy rates, although no significant correlation was found (Fig. 3). Lower ET_0_ values, below 3, resulted in relatively low efficacy rates. With further increases in ET_0_, there was a steady increase in the mean efficacy, as demonstrated by the trendline, yet with widely skewed predictions, indicated by the wider area around the trendline.

**Fig. 3.**
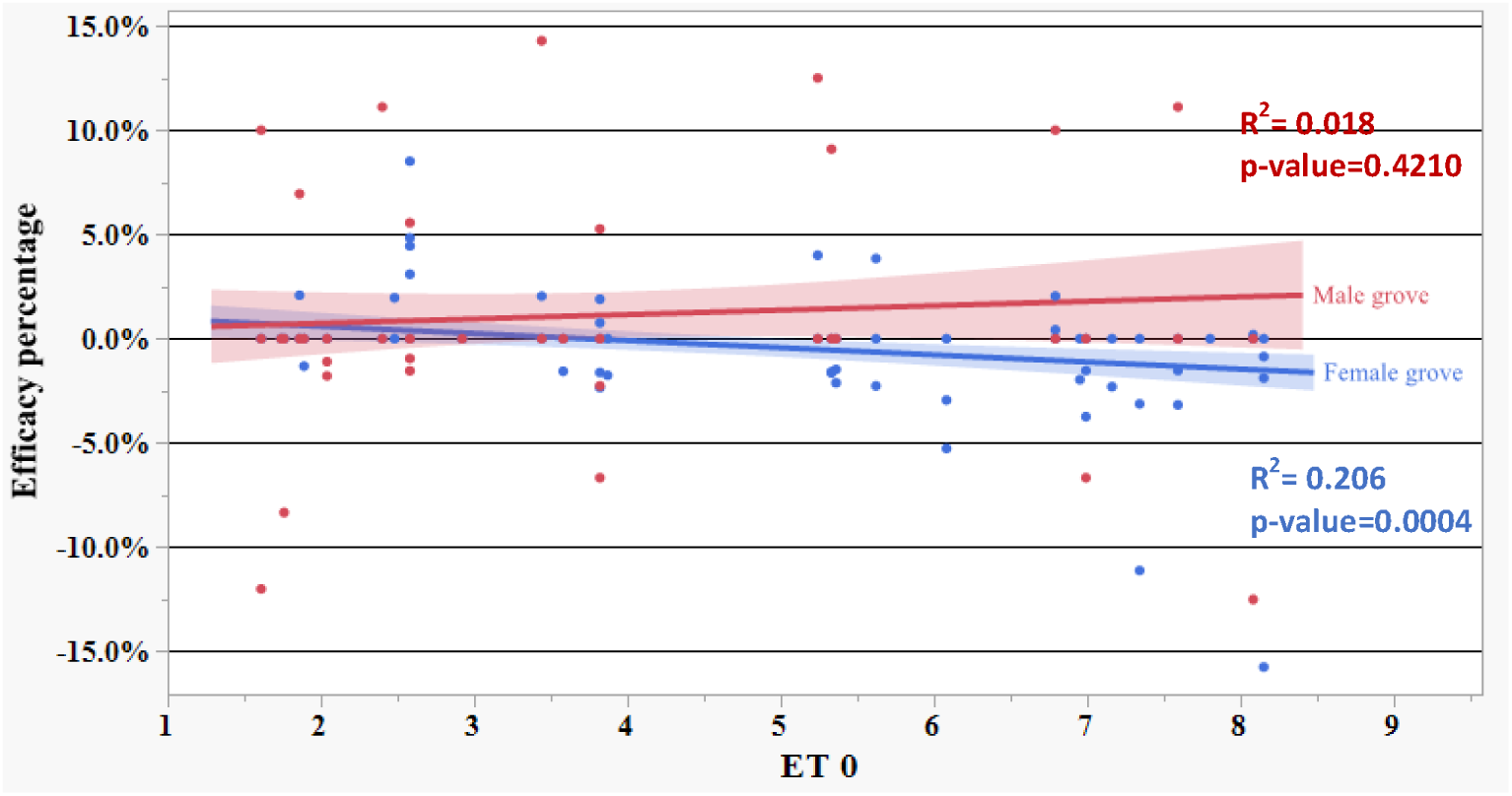
The relationship between climate variables (expressed as ET_0_ values) and the efficacy of EPF treatments compared to the control in the male (red) and female (blue) groves. Each dot represents an observation, and the linear regression trendline signifies the predicted efficacy according to ET_0_ values. A wider area around the trendline represents higher scattering of predictions. Positive efficacy percentages indicate a lower infestation rate for EPF treatments than the control, while negative efficacy percentages indicate a higher infestation rate.

### 3.4. EPF survival under different climate conditions

#### 3.4.1. Season, grove type and their combined effect on EPF survival

Whole-model analysis was performed to test the effects of season, agricultural practice, and the combined season x grove effect on EPF survival in the soil. Levene’s test for homogeneity of variances revealed unequal variances among the CFU loss per day results (F = 3.9952, *P* = 0.0039). Consequently, rank transformation was applied on this variable. Results did not show any significant combined season x grove effect on EPF survival (whole-model effect – season x grove: F = 0.2115, DF = 2, *P* = 0.8101). Similar survival rates were found in both groves across all seasons (whole-model effect – agricultural practice: F = 0.9069, DF = 1, *P* = 0.3454). In general, without considering the effect of grove variables, season had a significant effect on EPF survival (whole model effect – season: F = 6.0848, DF = 2, *P* = 0.0043). Post-hoc Tukey-HSD test revealed a significantly higher CFU loss in summer compared to autumn and spring. Spring and autumn EPF survival rates were approximately identical (Fig. 4).

**Fig. 4.**
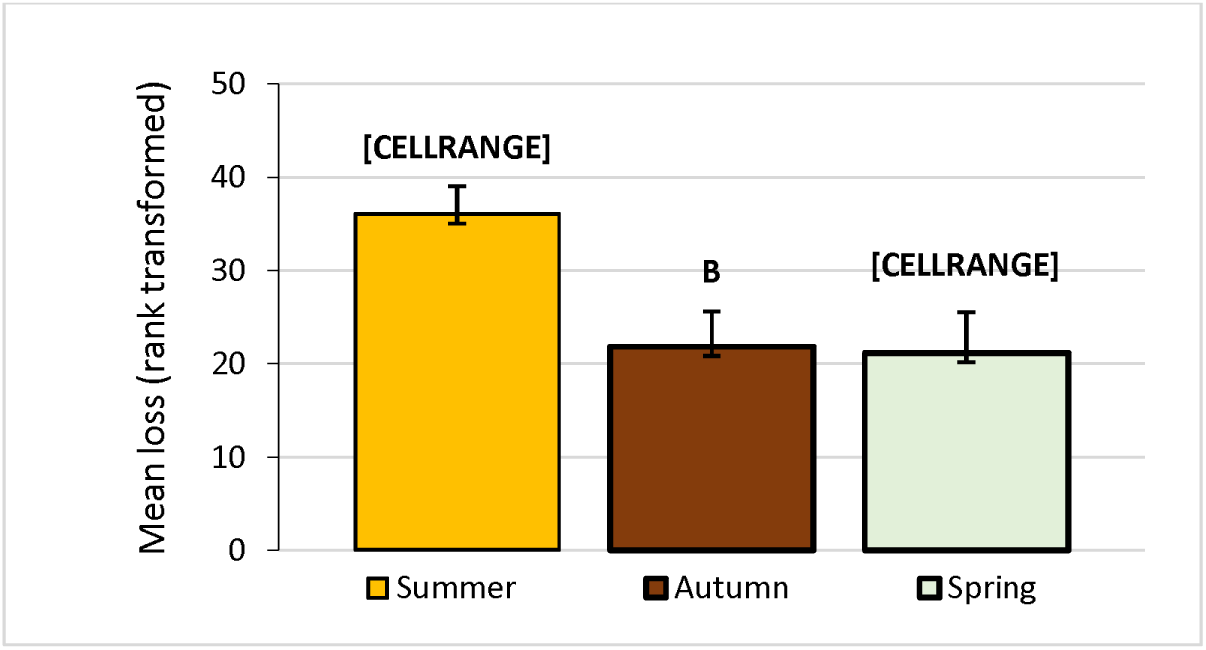
Mean CFU loss per day (ranked) as recorded in each season in both female and male groves in the Arava. Different uppercase letters indicate significant difference by Tukey-HSD test (α < 0.05).

#### 3.4.2. Season effect on EPF survival in each grove

Investigation of the impact of seasonal variations on CFU loss in the female grove revealed significant differences across different seasons. Initial analysis using Levene’s test indicated unequal variances among seasons (F = 7.5666, *P* = 0.0021). Consequently, Welch’s ANOVA was employed, revealing a significant effect of season on CFU survival (F = 6.3266, *P* = 0.0172). Accordingly, multiple-comparison t-tests were conducted between CFU loss per day for each pair of seasons, after verifying Levene’s equivalence of variance. A significant difference in survival rates was observed between summer and autumn (Levene’s test: F = 8.7233, *P* = 0.0066; t-test: t = 3.2017, *P* = 0.0082), However, comparisons between summer and spring did not reveal any significant difference in EPF survival (Levene’s test: F = 17.3730, *P* = 0.0004; t-test: t = 2.0426, *P* = 0.0910). Similarly, autumn and spring did not differ significantly in survival rates (Levene’s test: F = 0.5315, *P* = 0.4780; pooled t-test: t = 0.1044, *P* = 0.9183). Bonferroni correction was applied, setting α = 0.0167 to account for multiple comparisons (Table 10).

**Table 10.** Multiple comparisons between seasons for mean EPF differences in the female grove.

| (I)<br>Season 1 | (J)<br>Season 2 | Levene's<br>test $P$-value | Mean<br>difference<br>in CFU<br>loss<br>(I-J) | Std.<br>error | ANOVA test $P$-<br>value |
| --- | --- | --- | --- | --- | --- |
| Summer | Autumn | 0.0066 | 0.0201 | 0.0063 | **0.0082**<br># |
| Summer | Spring | 0.0004 | 0.0190 | 0.0093 | 0.0910<br># |
| Spring | Autumn | 0.4780 | 0.0018 | 0.01038 | 0.9183<br>@ |

Results in the male grove showed non-significant differences in EPF survival among the tested seasons (one-way ANOVA: F = 1.8820, *P* = 0.1783) Nevertheless, similar to the female grove, the largest CFU loss per day was observed in the summer.

#### 3.4.3. EPF survival variance among groves

Differences in EPF survival between male and female groves were examined in the summer, spring, winter, and autumn seasons. EPF survival rates did not differ between the groves in any season. Levene’s test was used to assess variance equality of the means. In the summer, survival rates were slightly, but not significantly, higher in the male group (Levene’s test: F = 25.5357, *P* > 0.0001; Welch’s t-test: t = 1.2603, *P* = 0.2331; Table 11). The autumn season showed very similar survival rates in both groves (Levene’s test: F = 0.0207, *P* = 0.8875; pooled t-test: t = 0.0449, *P* = 0.9647; Table 11). Survival differences were highest in the spring, when more EPF survived in the female grove, but results did not differ significantly (Levene’s test: F = 3.1399, *P* = 0.1068; pooled t-test: t = 0.5396, *P* = 0.6013; Table 11). Figure 5 illustrates the average daily CFU loss for each grove across different seasons and years. The data provide a descriptive comparison of CFU loss trends over time.

**Fig. 5.**
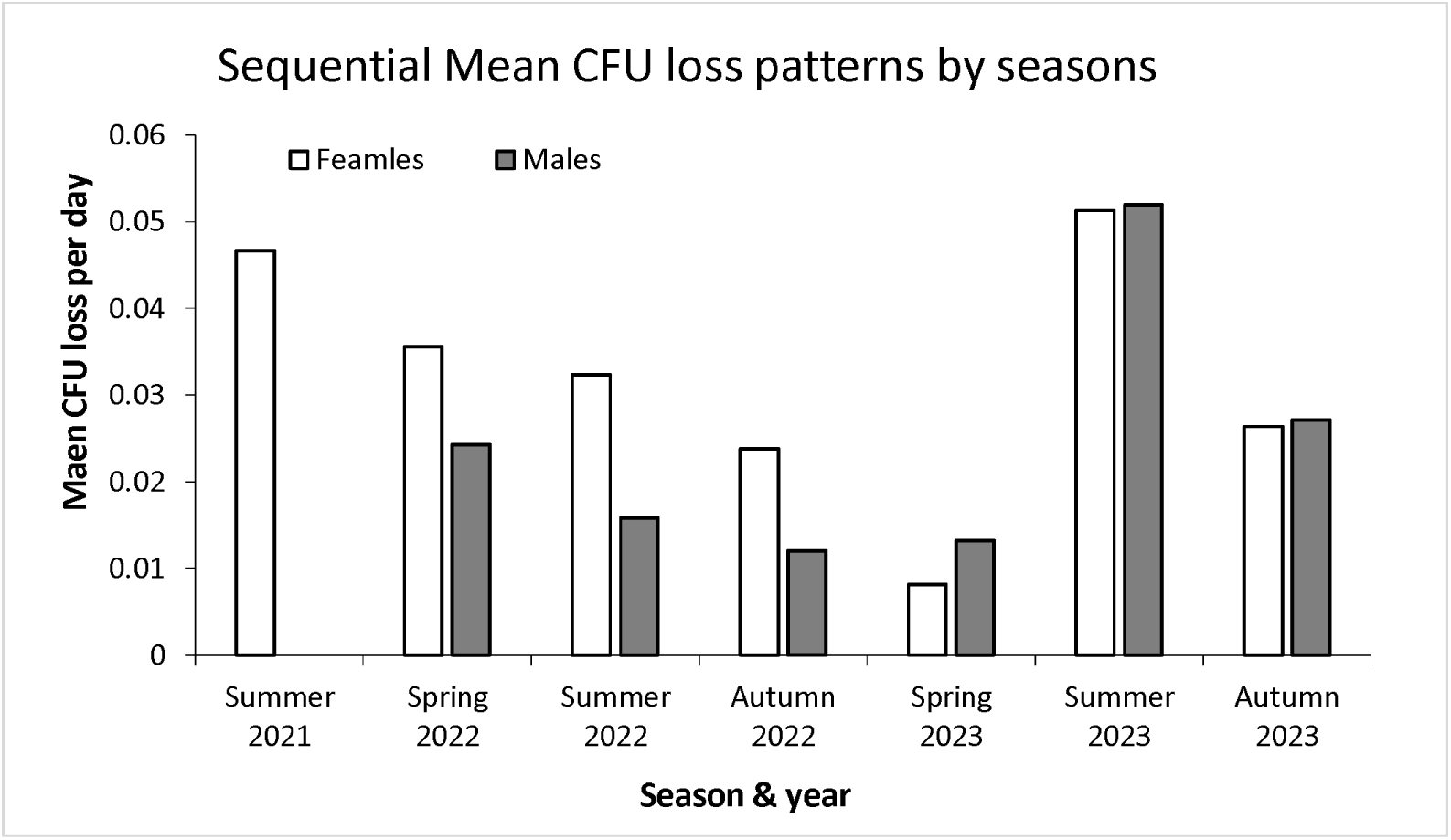
A descriptive comparison bar graph presenting mean CFU loss for female and male groves, ordered chronologically by seasons and years.

**Table 11.** Statistical test results for seasonal comparisons of survival rates between male and female groves.

| Season | DF | Levene's test<br>*P*-value | t-ratio | *P*-value |
| --- | --- | --- | --- | --- |
| Summer | 11.24 | >0.0001 | 1.2603 | 0.2331 @ |
| Autumn | 15 | 0.8875 | 0.0449 | 0.9647 # |
| Spring | 10 | 0.1068 | 0.5396 | 0.6013 # |

### 3.5. Association between EPF survival and ratio of healthy palms

The regression between the quantity of EPF surviving in the soil and the predicted percentage of healthy palms in EPF-treated and control groups for each experiment was tested, as described in section 2.7.4. Logistic fit test results demonstrated a significant positive association between CFU levels and palm tree health in the female grove (χ^2^ test, *p* = 0.0382). An upward trendline in the predicted percentage of healthy palms with increasing mean log CFU values. When the mean log CFU count at the end of the experiment was 0, indicating absence of EPF in the soil, the percentage of healthy palms was 88 and 95 in EPF and control groups, respectively (Fig. S1). The summed calculated probability for any palm with 0 CFU to be healthy was around 91%, as demonstrated by the trendline. As the mean log CFU reached 1 in the EPF treatment group, the predicted percentage of healthy palms rose to 92; 1.5–2.5 CFU/g soil, as found in the control groups, showed approximately 94% healthy palms. A further increase to 4 CFU/g soil in the EPF groups raised the predicted percentage of healthy palms in the female grove to almost 96.

A positive association between the quantity of EPF and palm health was also observed in the male grove, but without statistical significance (χ^2^ test, *P* = 0.4567). The regression trend in the male grove differed from that in the female grove (Fig. S2). When EPF was not found in the soil (0 CFU), approximately 84% healthy palms were predicted. An increase in CFU quantities from 0 to 1 increased the percentage of healthy palms to 90. As survival values rose to a mean log CFU of 2, there was a steady increase in predicted palm health, peaking at approximately 96% healthy palms. However, further increase to 3 and 4 CFUs corresponded to reductions in the predicted percentages of healthy palms to 90 and 86, respectively.

Logistic fit test results demonstrated a significant positive association between CFU levels and palm tree health when analyzing the results in both groves together (χ^2^ test, *P* = 0.0437), as demonstrated by the upward trendline in probability of healthy palms with increasing mean log CFU values (Fig. S3). When the mean log CFU count at the end of the experiment was 0, indicating absence of EPF in the soil, healthy palm probabilities ranged from 82% to 93% in the EPF and control groups, respectively. The summed calculated probability for any palm to be healthy with 0 CFU was around 91%, as demonstrated by the trendline. As the mean log CFU reached 1 in the EPF and control groups, the probabilities of healthy palms rose to 90% and 93%, respectively. In the control groups, soil with 1.5–2.5 CFU slightly increased the ratio of healthy palms, reaching its peak at approximately 94%. An increase in CFU among EPF-treated groups steadily raised the ratio of healthy palms, with 2, 3, and 4 CFU resulting in healthy palm probabilities of 92%, 93.5%, and 94.5%, respectively.

## 4. Discussion

EPF serve as effective tools against various crop pests, including RPW, but environmental factors affect their efficacy (Mascarin and Jaronski 2016; Ment et al. 2017; Qayyum et al. 2021; Sabbahi and Hock 2024). Our main objective was to study the efficacy of EPF application as a preventive treatment against RPW, EPF survival, and the relationship between the two in the face of biotic and abiotic factors in palm groves. Seasonal data on the survival and efficacy of EPF, along with their relationships in female and male date palms, is important for EPF assimilation in agricultural practice as a biologically friendly pesticide against RPW. Moreover, this information can further enhance the existing knowledge of EPF, improving their use as treatments in other crops and climatic areas.

Preventive EPF treatments against RPW showed the highest efficacy following autumn application in the female grove and spring application in the male grove. Different EPF treatment preparations did not affect the rate of protection efficacy. Regarding the climate data, ET_0_ values markedly affected the efficacy of EPF treatments in the female, but not male, grove. For example, mean ET_0_ values less than 4 correlated with effective EPF applications, whereas mean values exceeding 5 resulted in palm infestation rates that were higher than in the control. Regarding EPF survival rates, EPF propagule survival was lowest in the summer in both groves, with a marked difference in EPF survival between the summer and autumn in the female grove. In general, there was a significant positive correlation between the quantity of EPF propagules that survived in the soil and the percentage of healthy palms. However, when each grove was analyzed separately, this correlation was only observed in the female grove.

### 4.1. Seasonal variation of EPF survival

Seasonal changes significantly affected EPF survival, as reflected by the number of fungal propagules at the end of the experimental periods and, consequently, the calculated CFU loss per day. Summer season in the Arava Valley is extremely hot and dry (Ginat et al. 2011), conditions that do not favor *B. bassiana* persistence (Ment et al. 2017). Indeed, CFUs were not detected in most of the summer soil samples. Based on our calculations of CFU loss per day, it can be inferred that the death of EPF propagules likely occurred earlier than anticipated, resulting in a greater daily loss during the summer period. Climate conditions in the autumn and spring were optimal or near-optimal for EPF survival. These results correspond with a previous study in Egypt by Hussein et al. (2010), who found that *B. bassiana* is more abundant during the spring and autumn than the summer and winter; and that the factors with the strongest effect on *B. bassiana* prevalence in the soil are maximum and minimum temperatures and minimum humidity levels. Comparison of the number of fungal propagules at the end of the experiment among the groves revealed higher persistence mostly in the male grove. However, the effect of the grove itself, or in combination with the effect of season, on EPF survival was not significant. This observation suggests that the fungal propagules have a limited protective effect in the palm canopy; alternatively, it can be attributed to the smaller amount of experimental data from the male grove compared to the female grove.

### 4.2. Efficacy of preventive EPF application in different seasons

Climate conditions have a strong effect on EPF survival, pathogenicity, and therefore, efficacy (reviewed in Ment et al. 2017). RPW activity was previously found to be lacking in the summer season in the Arava Valley, with no palm infestation, due to the extremely high temperatures in the area (Mendel et al. 2024b). In the present study, RPW infestation rate was compared among EPF-treated date palms and untreated control palms over the years among seasons. Autumn EPF application resulted in the highest protection efficacy among all tested seasons. During the autumn, mean ET_0_ rates were approximately 3, and the season was characterized by low radiation flux and a mean temperature of 23.6 °C. These conditions are expected to result in high rates of *B. bassiana* survival, infectivity, and sporulation (Ment et al. 2017; Zaman et al. 2020). In general, favorable climatic parameters for EPF development were accompanied by significant EPF survival and palm protection.

Spring trials gave ambiguous results. In general, EPF application showed no advantage over the control group in protecting palms from RPW. A comparison of spring and autumn climate factors revealed higher ET_0_ values, characterized by very similar temperatures in both seasons but lower RH and higher UV radiation in the spring. Our assumption is that these differences decreased EPF survival and pathogenicity, resulting in lower EPF efficacy in this season. Analysis of the results in each grove revealed efficient protection in the male grove but not in the female grove. These variations can be explained by the UV-protective shield effect of the shorter palms and lower canopies in the male grove. Another potential explanation is that the application of ammonium nitrate and urea in the male grove stimulated EPF growth and sporulation (Lingg and Donaldson 1981; Cojocaru and Lumînare 2021), thereby compensating for the loss of fungal propagules during the trials.

Summer trials were characterized by high temperatures, low RH, and high UV radiation, leading to decreased EPF efficacy at protecting palms from RPW. Abiotic factors in the summer months do not favor *B. bassiana*. The mean recorded ET_0_ value was 7.35, with a mean temperature exceeding 30 °C, while RH was <50% in all summer months. Field trials conducted by Tehri et al. (2015) similarly demonstrated the ineffectiveness of *B. bassiana* in controlling *Tetranychus urticae* populations when applied during hot and dry crop seasons. However, unexpected and surprising were the significantly higher rates of RPW-infested palms among EPF-treated vs. control trees. Looking at each grove separately, this phenomenon was only found in the female grove. We failed to find similar effects in previous experiments. Although this experimental setup cannot be replicated to gain further data for analysis, another speculation may involve the effect of thermal stress on EPF and its volatile organic compounds, which manipulate arthropods’ behavior (Liu et al. 2008; Ramírez-Ordorica et al. 2022; Ma et al. 2024).

### 4.3. Effect of EPF preparations on seasonal protection efficacy

Applications of different EPF treatments, which included different *B. bassiana* strains, surfactants, or a double dose, had no significant effect on the protection rate against RPW infestation in either grove. The Velifer application during the winter and spring showed similar or non-significantly higher protection rates compared to Botanigard. This solid *B. bassiana* formulation has been previously shown to control populations of RPW when dusted around the crown, stems, and petioles of Canary palms (Güerri-Agulló et al. 2012) and Washingtonia palms (Ment et al. 2023). Despite its promising potential, Velifer was unavailable for further trials. Addition of the surfactant L77 to Botanigard did not significantly impact its effectiveness. The most surprising result was the lack of impact of a double-dose application of Botanigard, which had been previously proven to positively correlate with efficacy rates (Kaur et al. 2011). However, caution should be exercised when drawing conclusions about the differences among treatments. Except for the Botanigard treatments, EPF treatments were applied with a low number of repetitions and relatively small sample sizes, which could impact the validity of the results. Analyzing the seasonal variations of each EPF preparation against the control group revealed significant differences in treatment efficacy only in the female grove for Botanigard + L77 treatments, with autumn applications demonstrating superior protection compared to summer applications. Again, it is possible that the relatively low sample size in this group was not sufficient to achieve statistical significance.

### 4.4. Association between EPF CFU level and palm-protection rates

Summer trials were characterized by a significant loss of EPF propagules compared to the autumn trials. These two seasons also showed opposite trends in RPW-prevention efficacy, with significant protection in autumn. A similar correlation between *B. bassiana* persistence in the soil and its conidia’s infectivity of the lettuce aphid *Nasonovia ribisnigri* (Mosley) was found in semi field experiments (Shrestha et al. 2015).

Relying on a previous study in which the survival of EPF propagules was found to be similar between the trunk and soil of treated palms (Livne 2024; Ment unpublished results), in the current study, EPF persistence was evaluated for soil samples only. Therefore, we decided to use EPF CFUs in the soil as a measure of fungal persistence. Moreover, as the protection mechanism following EPF application relies on fungal propagule administration to the egg-laying hole created by the RPW female (Matveev et al. 2023), fungal persistence is expected to correlate with palm health. Indeed, results indicated that EPF CFU/g soil surrounding treated palms is positively correlated with palm health. Although not statistically significant, the male grove regression pattern demonstrated a positive correlation between CFU rates and a preventive effect in the control palms. Conversely, EPF-treated palms showed a decrease in efficacy with increasing CFU rates. Regarding the control palms, we mentioned that EPF-treated and control groups were arranged in clusters, with at least 15 m between groups. Although the recorded rates of log 1 CFU, representing 10 CFU/g soil, are relatively low in terms of EPF rates, they were retrieved approximately 3 months after application from untreated soil. The increase in CFU levels, accompanied by lower infestation rates, suggests that EPF conidia can disperse and become established throughout the grove, affecting untreated palms as well. The reason for the negative regression in treated palms, however, remains unclear. We can hypothesize that in the male grove, fungal persistence is relatively high due to the UV protection provided by lower canopies. Nevertheless, pathogenicity and virulence are influenced by temperature and humidity, and thus, they may not be correlated with spore survival.

### 4.5. Research limitations and future prospects

Our research focused on the effects of each season’s environmental conditions on the survival of the EPF *B. bassiana* and its efficacy in preventing RPW infestation in male and female commercial date palm groves. Our study included 250 female palms and 56 male palms, some of which were excluded due to our experimental design. In most cases, these sample sizes were not large enough to provide significant results. The RPW typically targets a relatively small percentage of palms, further highlighting this limitation. In addition, it is important to note that the climate conditions recorded during our experiments were obtained from a weather station situated in an open field and as such, may not fully depict the actual conditions found under the date palms. Livne (2024) observed significantly reduced solar radiation levels and slight temperature variations within a grove of date palms compared to a nearby weather station in an open field. Moreover, morphological differences between the palms, deriving from palm variety and growth characteristics, can alter the frond and leaf numbers and lengths (Zaid 2002) which, in turn, can decrease the UV radiation and temperature in the palm trunk and its surroundings. Another limitation in our field test was the multicollinearity of the climatic factors temperature, UV, and RH. Because it is not possible to isolate and assess the influence of each factor on its own, we considered ET_0_ values, which are commonly used as a general representative index of these abiotic factors. Furthermore, our comparison of CFU loss/day rates assumed that EPF has a linear declining rate, which does not describe EPF loss comprehensively. In addition, a count of 0 CFU at the end of a season could suggest an even higher rate of EPF propagule loss. Soil samples taken from the male grove in December 2021, as well as several samples from both groves in different seasons were lost and were therefore subtracted from the data analysis. We suggest that future investigations include the examination of a larger number of palms, which can yield more significant results under varied environmental conditions. Another study might test EPF survival in the trunk and/or in the soil of the palms on a monthly basis to characterize seasonal EPF survival patterns more accurately in the different seasons and groves. Assessment of pathogenicity and virulence under seasonal variations could expose another important element. Studies that include different EPF species, strains, and formulations can also enrich the existing knowledge and promote the use of EPF as a biologically friendly insecticide.

## Author contributions

DM, OZ and SM conceived and designed research. OZ and SM conducted experiments. DM, OZ, and SM analyzed data. DM and OZ wrote the manuscript. DM recruited funds. All authors read and approved the manuscript.

## Acknowledgements

We are grateful to the farmers of Moshav Idan in northern Arava, Itamar Haviv and Moti Arnon for contributing the date palm groves for the experiments and for the invaluable assistance in the different applications. We acknowledge the professional statistics consultancy provided by Dr. Hillary Voet. We are in debt for the assistance provided by Svetlana Dobrinin and Dr. Shai Daniel of the Ministry of Agriculture Extension Services during the research. We acknowledge Rimi for providing invaluable assistance and providing Botanigard throughout the research. We acknowledge the assistance provided by Biobee for providing Velifer. We appreciate the technical support provided by Agrint in the maintenance and data acquisition of the seismic sensors.

## Funding

The plant protection & marketing board for an award to OZ for the years 2022 and 2024. ICA In Israel (project no. 279/21) and Plant board (project no. 132234422) awarded to DM.

## Supplementary Information

**Table S1.** Monthly recordings of mean, maximum, and minimum values of temperature, relative humidity, solar radiation, and mean evapotranspiration rates at the study area in the Arava.

| Month | Temperature<br>(°C) |  |  | Relative humidity<br>(%) |  |  | Radiation<br>(W/m<sup>2</sup>) |  |  | ET<sub>0</sub><br>(mm/day) |
| --- | --- | --- | --- | --- | --- | --- | --- | --- | --- | --- |
|  | Mean | Max | Min | Mean | Max | Min | Mean | Max | Min | Mean |
| Jun 21 | 31 | 43.6 | 21.9 | 38 | 68 | 12 | 345 | 1035 | 0 | 7.8 |
| Jul 21 | 34.1 | 44.7 | 25.9 | 36 | 68 | 9 | 331 | 1024 | 0 | 8.15 |
| Aug 21 | 34.5 | 46.2 | 26.1 | 36 | 68 | 6 | 305 | 988 | 0 | 6.95 |
| Sep 21 | 30.8 | 40.9 | 19.9 | 46 | 83 | 16 | 244 | 921 | 0 | 5.24 |
| Oct 21 | 27.5 | 36.6 | 19 | 47 | 76 | 19 | 175 | 840 | 0 | 3.44 |
| Nov 21 | 22.2 | 33.7 | 12.9 | 48 | 86 | 14 | 135 | 698 | 0 | 1.86 |
| Dec 21 | 17.2 | 26.9 | 8.3 | 49 | 83 | 19 | 107 | 617 | 0 | 1.61 |
| **Jan 22** | 13.8 | 24.5 | 3.1 | 57 | 92 | 22 | 120 | 725 | 0 | 1.76 |
| **Feb 22** | 16.4 | 26.2 | 7.6 | 54 | 90 | 15 | 150 | 825 | 0 | 2.4 |
| **Mar 22** | 17.3 | 32.7 | 6.5 | 44 | 80 | 9 | 198 | 962 | 0 | 3.58 |
| **Apr 22** | 25.5 | 42.4 | 14.2 | 34 | 75 | 5 | 261 | 1035 | 0 | 5.33 |
| **May 22** | 28.3 | 41.9 | 16.3 | 33 | 72 | 3 | 310 | 1031 | 0 | 6.79 |
| **Jun 22** | 31.8 | 43.9 | 23 | 36 | 71 | 9 | 338 | 1024 | 0 | 7.59 |
| **Jul 22** | 33.3 | 43.5 | 23.6 | 37 | 67 | 10 | 334 | 1009 | 0 | 8.081 |
| **Aug 22** | 33.8 | 43.5 | 25.3 | 43 | 68 | 13 | 289 | 952 | 0 | 6.99 |
| **Sep 22** | 31.5 | 40.2 | 22.3 | 45 | 74 | 15 | 236 | 888 | 0 | 5.36 |
| **Oct 22** | 28 | 41 | 20.2 | 50 | 90 | 17 | 180 | 782 | 0 | 3.82 |
| **Nov 22** | 22.3 | 30.6 | 12.6 | 53 | 90 | 21 | 151 | 712 | 0 | 2.58 |
| **Dec 22** | 17.5 | 27.1 | 8.8 | 63 | 94 | 27 | 133 | 612 | 0 | 1.74 |
| **Jan 23** | 16.3 | 29.8 | 6.8 | 58 | 96 | 21 | 146 | 667 | 0 | 2.04 |
| **Feb 23** | 16.3 | 33.1 | 7.6 | 50 | 86 | 12 | 185 | 839 | 0 | 2.921 |
| **Mar 23** | 20.4 | 32.2 | 11 | 48 | 92 | 10 | 219 | 963 | 0 | 3.87 |
| **Apr 23** | 23.4 | 37.2 | 13.1 | 39 | 69 | 10 | N.A | 1037 | 0 | 2.48 |
| **May 23** | 27.4 | 42.1 | 18.6 | 35 | 82 | 6 | N.A | 1026 | 0 | 6.08 |
| **Jun 23** | 31 | 44 | 21.5 | 38 | 69 | 10 | 319 | 1026 | 0 | 7.34 |
| **Jul 23** | 34.8 | 45.9 | 25.1 | 34 | 67 | 10 | 332 | 1014 | 0 | 8.15 |
| **Aug 23** | 33.9 | 43.6 | 25.5 | 45 | 69 | 9 | 288 | 994 | 0 | 7.16 |
| **Sep 23** | 31.9 | 44 | 22.9 | 43 | 70 | 13 | 241 | 878 | 0 | 5.62 |
| **Oct 23** | 28.6 | 36.9 | 20.8 | 48 | 78 | 20 | 177 | 822 | 0 | 3.82 |
| **Nov 23** | 23.7 | 35.6 | 12.4 | 52 | 87 | 20 | 153 | 676 | 0 | 2.58 |
| **Dec 23** | 18.8 | 29.2 | 11.1 | 59 | 93 | 28 | 130 | 640 | 0 | 1.89 |

**Fig. S1.**
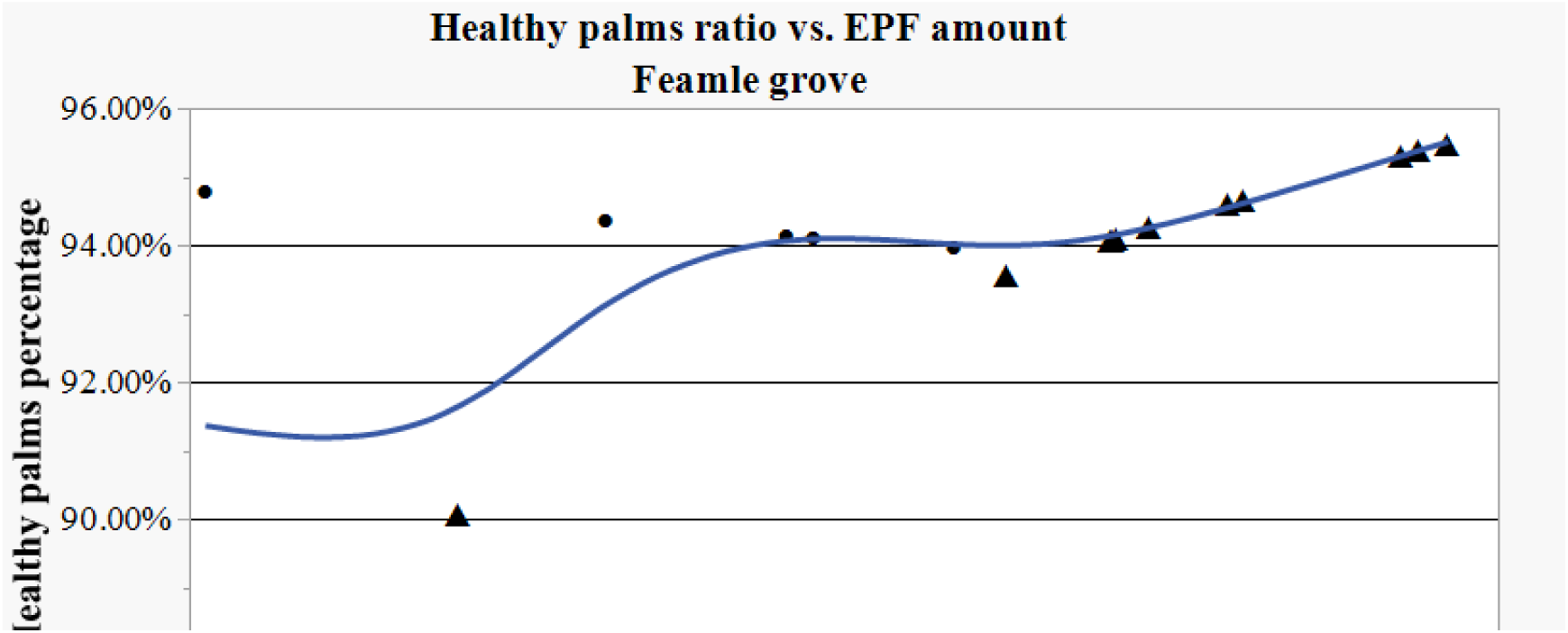
Relationship between soil EPF levels and palm health in the female grove. ● - control group predictions; ▴ - EPF group predictions

**Fig. S2.**
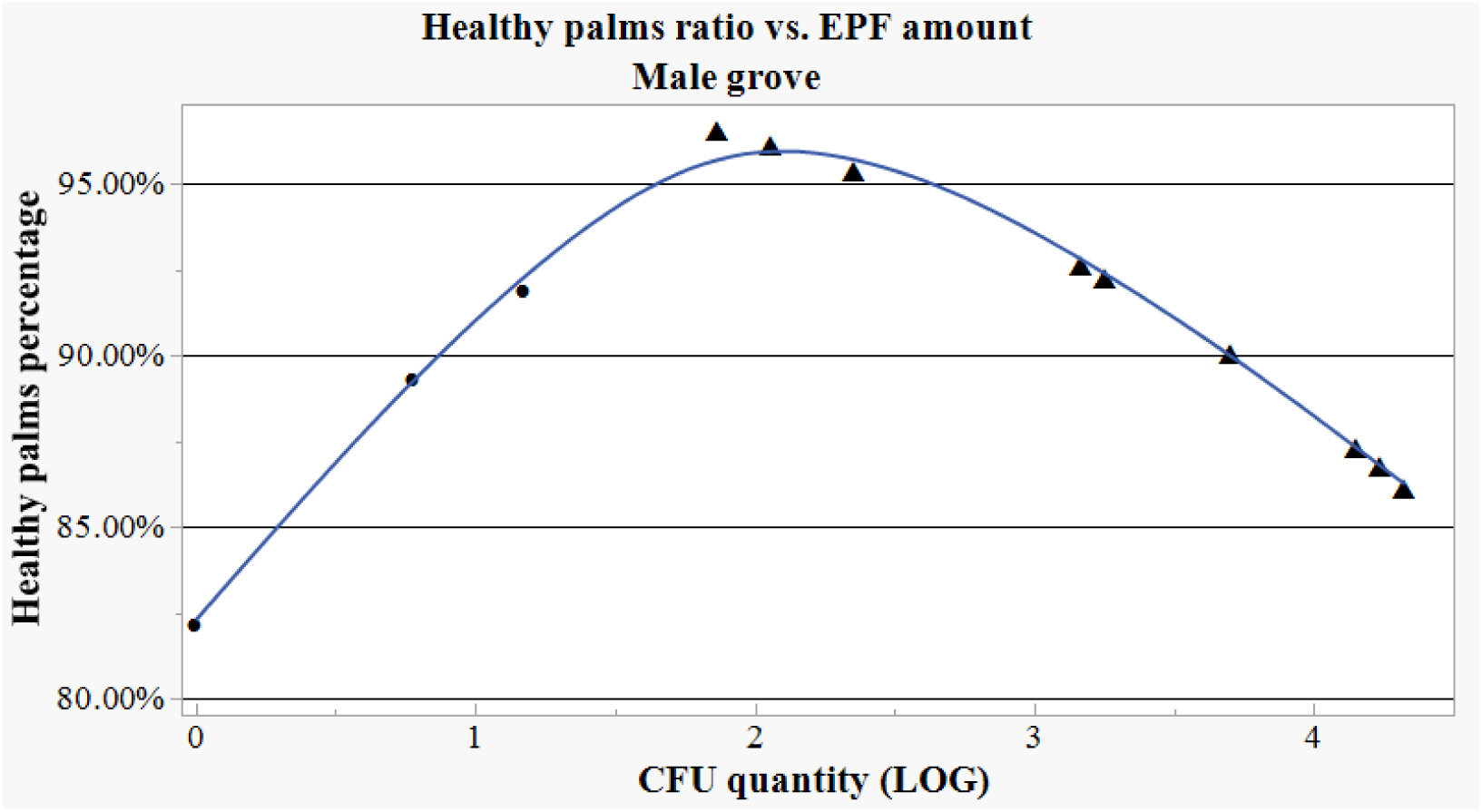
Relationship between soil EPF levels and palm health in the male grove. ● – control group predictions;▴ - EPF group predictions

**Fig. S3.**
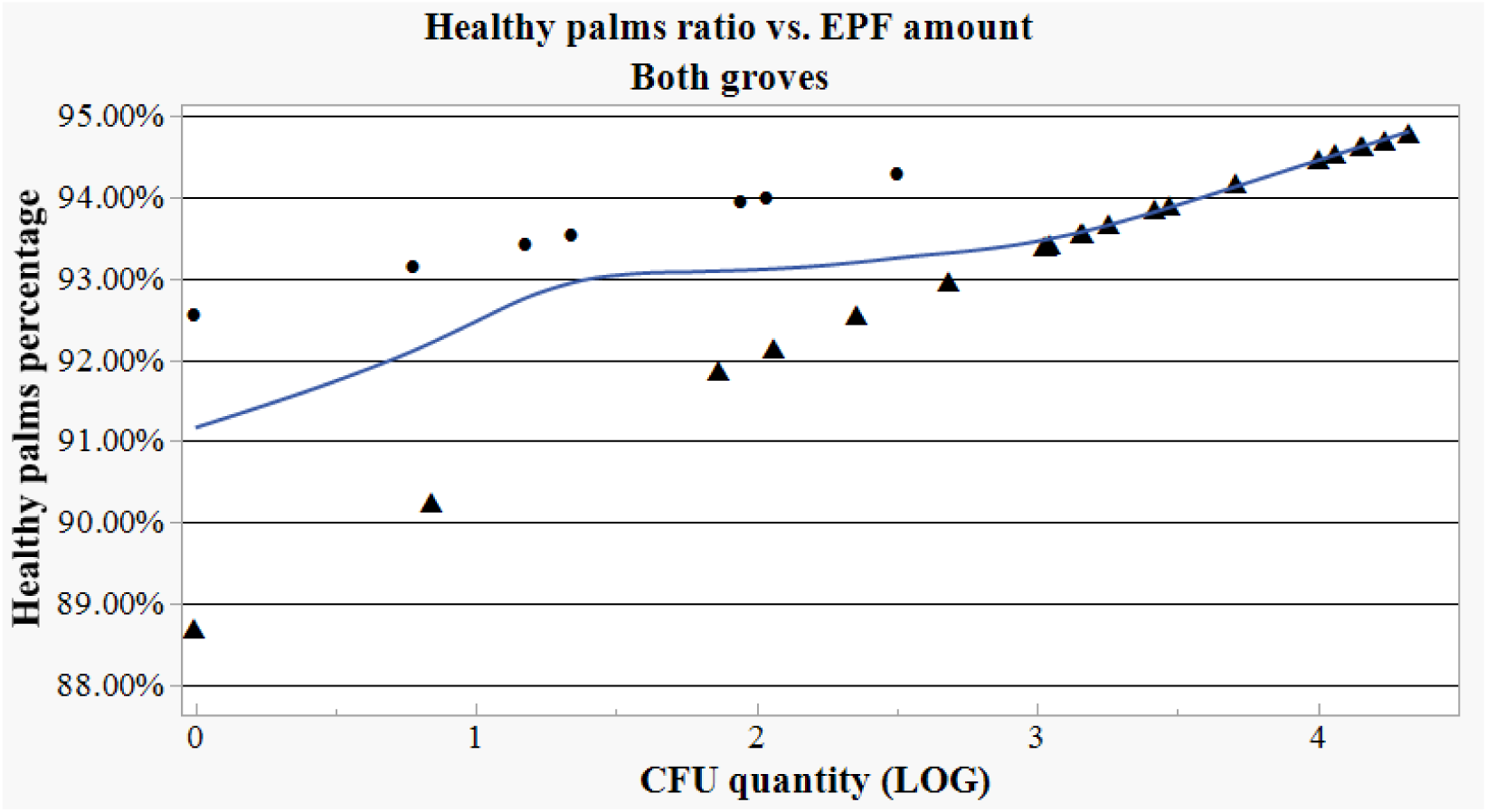
Relationship between soil EPF levels and palm health in both male and female groves. ● - control group predictions; ▲ - EPF group predictions.

## Notes

### Competing Interest Statement

The authors have declared no competing interest.

### Summary of Updates

The current version of the manuscript has undergone language editing bu a qualified English scientific editor. Part of the results were moved into the supplementary and the result part was thus shortened.

